# Neutralization of SARS-CoV-2 Omicron BA.4/BA.5 subvariant by a booster dose of bivalent adjuvanted subunit vaccine containing Omicron BA.4/BA.5 and BA.1 subvariants

**DOI:** 10.1101/2022.10.07.511263

**Authors:** Tsun-Yung Kuo, Chia En Lien, Yi-Jiun Lin, Meei-Yun Lin, Luke Tzu-Chi Liu, Chung-Chin Wu, Wei-Hsuan Tang, Charles Chen

## Abstract

The dominance of SARS-CoV-2 variants of concern (VoC), such as the Omicron subvariants, is a threat to the current vaccination scheme due to increased resistance to immune neutralization and greater transmissibility. To develop the next generation of prefusion SARS-CoV-2 spike protein (S-2P) subunit vaccine adjuvanted with CpG1018 and aluminum hydroxide, mice immunized with two doses of the adjuvanted ancestral Wuhan strain (W) followed by the third dose of the W or Omicron variants (BA.1 or BA.4/BA.5) S-2P, or a combination of the above bivalent S-2Ps. Antisera from mice were tested against pseudovirus neutralization assay of ancestral SARS-CoV-2 (WT) and Omicron BA.4/BA.5 subvariant. Boosting with bivalent mixture of Omicron BA.4/BA.5 and W S-2P achieved the highest neutralizing antibody titers against BA.4/BA.5 subvariant pseudovirus compared to other types of S-2P as boosters.

## Introduction

The development and production of Coronavirus Disease 2019 (COVID-19) vaccines have altered the course of this disruptive pandemic, averting millions of deaths globally. The first year of COVID-19 vaccination has prevented approximately 19.8 million deaths from the disease in 185 countries and territories [1]. However, mutations in the severe acute respiratory syndrome coronavirus 2 (SARS-CoV-2) have resulted in the emergence of viral strains that are more lethal and resistant to the protection offered by vaccines rendering the originally developed vaccines less effective against these new strains. As a result, the primary series, and even booster doses, of existing vaccines have been found to induce less neutralization against the Omicron variant and its subvariants, including, and most significantly, the currently dominant BA.4 and BA.5 subvariants [2, 3]. Given the situation, second-generation vaccines must be developed. To date, vaccine companies have begun developing second-generation vaccines, particularly, Omicron-adapted candidates. Among these are the mRNA vaccines developed by Moderna and Pfizer-BioNTech, which are bivalent vaccine based on the original SARS-CoV-2 and the Omicron BA.4/BA.5 subvariants for individuals aged 18 years and older [4, 5]. Aside from these two manufacturers, companies such as Novavax and Sinocelltech were conducting phase III clinical trials for Omicron-based and bivalent variant subunit vaccines [6, 7].

MVC-COV1901 is a subunit vaccine against COVID-19 based on the stable prefusion spike protein S2-P of the ancestral SARS-CoV-2 and adjuvanted with the CpG 1018 adjuvant and aluminum hydroxide [8]. A previous study has shown that three doses of MVC-COV1901 can increase neutralization against the Omicron variant pseudovirus [9]. In a preclinical study, the vaccine has also demonstrated the ability of the Beta variant version of the S2-P to offer cross-protection against different variants, including the Omicron variant [10]. When used as a booster in a phase I clinical trial, the Beta version of MVC-COV1901 (MVC-COV1901-Beta) showed a favorable safety profile and broad immune response with cross-immunity with Omicron and the pseudovirus neutralization assay results indicated neutralizing antibody titers are higher than the ones induced by a booster dose of the original MVC-COV1901 [11]. Considering the continuous evolution of the virus, developing an Omicron-based vaccine is a crucial next step in the process. In this study, we evaluated the immunogenicity of Omicron BA.1 or BA.4/BA.5 versions of adjuvanted S-2P as boosters after two doses of original adjuvanted S-2P against BA.4/BA.5 pseudovirus neutralization in a mouse model.

## Methods

Female BALB/c mice aged 6-9 weeks at study initiation were obtained from BioLASCO Taiwan Co., Ltd. (Yilan County, Taiwan). Animal immunizations were conducted in the Testing Facility for Biological Safety, TFBS Bioscience Inc., Taiwan. All procedures described in this study involving animals were performed in a manner to avoid or minimize discomfort, distress, or pain to the animals and were carried out in compliance with the ARRIVE guidelines (https://arriveguidelines.org/). All animal work in the current study was reviewed and approved by the Institutional Animal Care and Use Committee (IACUC) with animal study protocol approval number TFBS2020-006.

Intramuscular administration of the vaccine was performed by injection in the quadriceps femoris muscle of the left and right legs of each hamster (50 μL each leg for a total of 100 μL per dose). Three immunizations were performed on days 1, 22, and 43. Hamsters were grouped into six groups (n = 5 per group), as shown in Table 1, and immunized with the following regimens:

**Table 1.**
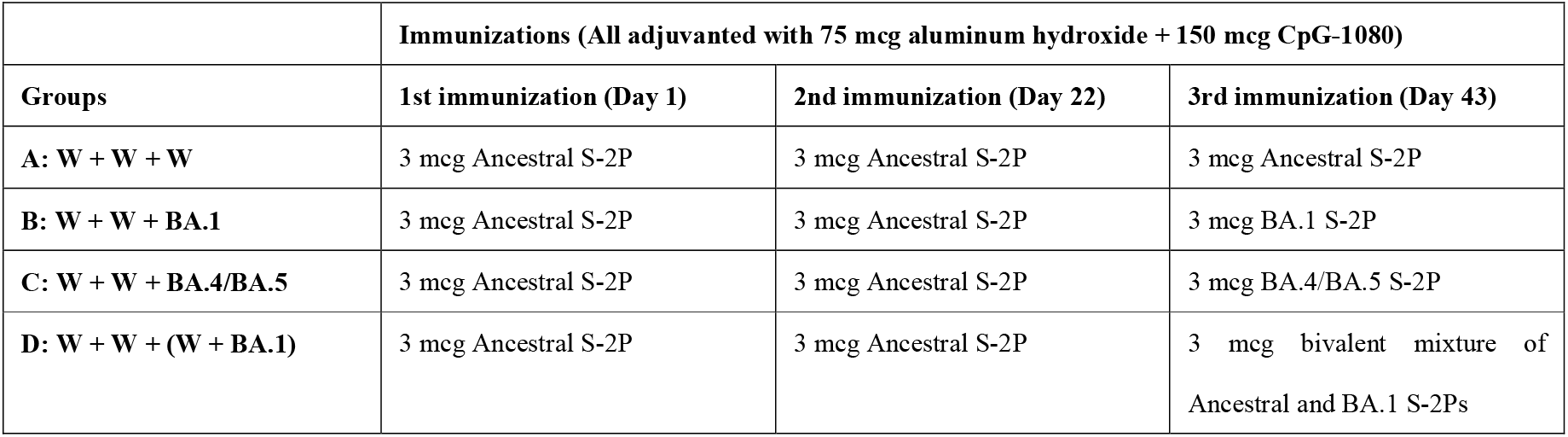
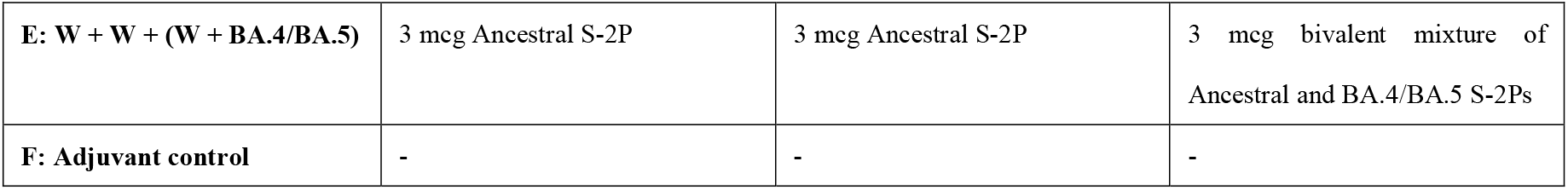

Serum samples were harvested on days 36 and 57 (i.e. 2 weeks after the second and third immunizations, respectively) and were subjected to pseudovirus neutralization assay.

Pseudotyped lentivirus with spike proteins of Omicron BA.1 or BA.4/BA.5 (with BA.4 and BA.5, both possess identical spike protein sequences) variant was used in pseudovirus neutralization assay conducted as reported previously [8]. Results were reported as 90% inhibition dilution (ID_90_) geometric mean titer (GMT) with 95% confidence intervals (95% CI), as shown in Figure 1. The mutations for the Omicron variant (BA.1 and BA.4/BA.5) used in the spike sequence for pseudovirus and S-2P construction were based on the WHO VoC profiles of spike amino acid changes [12].

**Figure 1.**
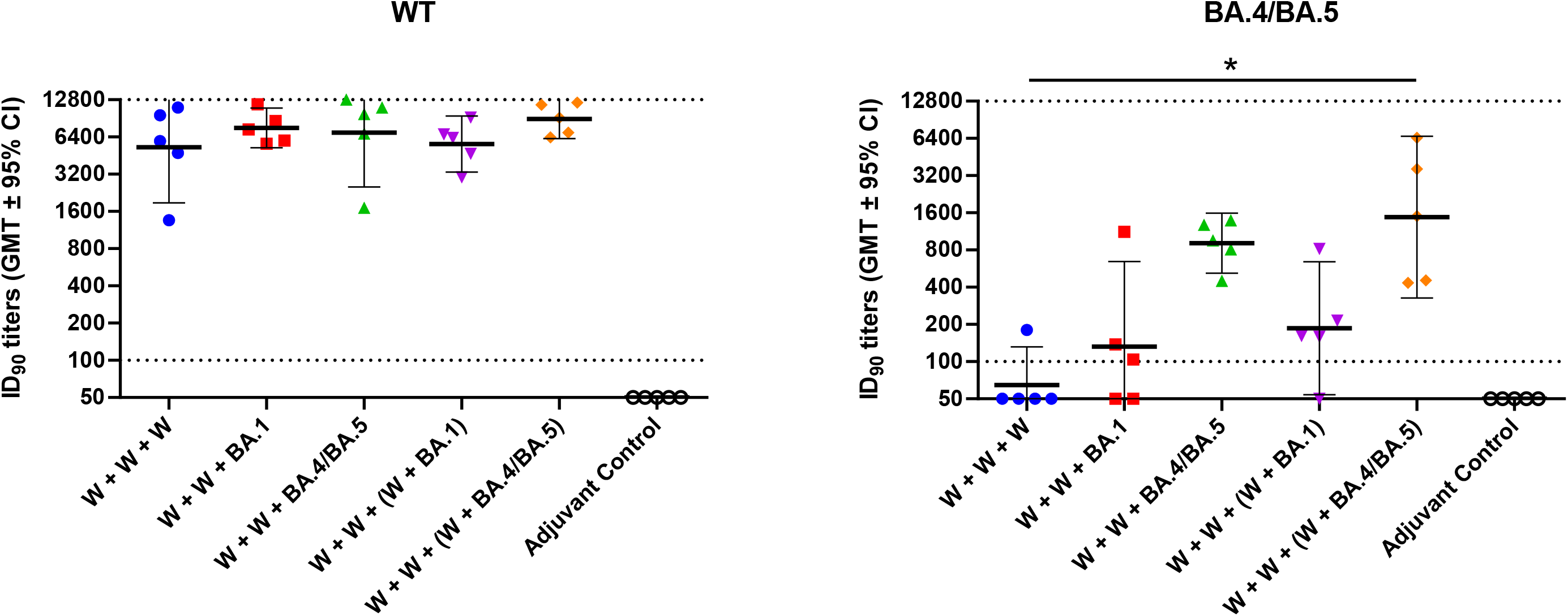
Pseudovirus neutralisation assay of pseudovirus with spike proteins of the SARS-CoV-2 A). ancestral Wuhan strain (WT) or B). BA.4/BA.5 with mice serum samples from two weeks after the third immunization. BALB/c mice (n = 5 per group) were immunized with combinations of adjuvanted S-2P derived from the ancestral SAR-CoV-2 (W) or Omicron (BA.1 or BA.4/BA.5) subvariants as indicated on the x-axis. Antisera were harvested two weeks after the third dose and subjected to neutralization assay with pseudovirus expressing SARS-CoV-2 ancestral strain (WT) or BA.4/BA.5 spike proteins. Shaped dots show individual titer values, horizontal bars represent GMTs and error bars represent 95% confidence intervals of the 90% inhibition dilution (ID_90_). Dotted lines indicate the starting dilution (100; lower dotted line) and the final dilution (12800; upper dotted line) for the assay. Kruskal-Wallis with corrected Dunn’s multiple comparisons test was used to calculate significance. * = p < 0.05, ** = p < 0.01, *** = p < 0.001, **** = p < 0.0001

Prism 6.01 (GraphPad, San Diego, CA) was used for statistical analysis. Kruskal-Wallis with corrected Dunn’s multiple comparisons test was used to calculate significance. * = p < 0.05, ** = p < 0.01, *** = p < 0.001, **** = p < 0.0001

## Results

A booster dose of adjuvanted S-2P derived either from the ancestral SARS-CoV-2 (W) or Omicron (BA.1 or BA.4/BA.5) variant or bivalent mixtures of the above was administered to mice after two doses of adjuvanted W S-2P. The immunogenicity was assessed two weeks after the third immunization by pseudovirus neutralization assay with ancestral Wuhan strain SARS-CoV-2 (WT) and Omicron BA.4/BA.5 subvariant. As shown in Figure 1A, all combinations of the S-2Ps induce equally high neutralization antibody titers against the WT. The group with two doses of W S-2P boosted with the bivalent W + BA.4/BA.5 S-2Ps achieved the highest numerical GMT (8946 [95% CI: 6211 – 12886]), while boosting with BA.1 and BA.4/BA.5 resulted in GMT of 7581 [5246 – 10955] and 6942 [2517 – 19145], respectively (Figure 1A). In comparison, three doses of W S-2P resulted in the lowest GMT out of all treatment groups (5274 [1877 – 14824]), although the GMTs are not significantly different among the treatment groups (p > 0.05).

In Figure 1B, three doses of W S-2P were ineffective against the BA.4/BA.5 pseudovirus with the GMT below the limit of detection (64.6 [31.7 – 131.5]). Boosting with either BA.1 or a bivalent mixture of W and BA.1 S-2Ps after two doses of W S-2P only slightly increased the GMTs (132.2 [27.1 – 645.8] and 186.3 [54 – 642.7], respectively). A booster dose of BA.4/BA.5 S-2P after two doses of W S-2P had a higher increase in GMT to a value of 907.9 [519.4 – 1587]. The only treatment that resulted in significantly higher neutralization titer (p < 0.05) against the BA.4/BA.5 pseudovirus was the bivalent mixture of W and BA.4/BA.5 S-2Ps with a GMT of 1473 [326.4 – 6645] (Figure 1B).

## Discussion

Two doses of W S-2P with a booster dose of a bivalent mixture of W and BA.4/BA.5 S-2Ps induced the highest neutralizing antibody titer against both pseudoviruses based on the ancestral SARS-CoV-2 and the Omicron BA.4/BA.5 subvariant. The results suggested that boosting with BA.4/BA.5 S-2P or a mixture of W and BA.4/BA.5 S-2Ps can greatly increase the neutralizing titer against the BA.4/BA.5 subvariant compared to boosting with W or BA.1 S-2Ps. This may be due to immune refocusing and the expansion/selection of cross-reactive neutralizing antibodies recognizing more conserved regions of the S-2P after boosting with the variant-based S-2P [13]. Another subunit vaccine currently in use, NVX-CoV2373 by Novavax, has been shown to induce increased neutralizing antibody titers against the BA.1 and BA.4/BA.5 after three homologous doses [14]. Novavax has also initiated a phase III trial using booster doses based on the BA.1 and BA.5 subvariants, and a bivalent mixture of the original NVX-CoV2373 and one of the above subvariants [15]. Since the variant-based and bivalent COVID-19 vaccines have only been approved recently, how the effectiveness against the Omicron subvariants in the real world is yet unclear. However, in a mouse challenge study, boosting with Omicron BA.1-based mRNA vaccine protected mice from Omicron BA.1 infection; thus, this result could translate to protection from the Omicron variant in humans [16]. The currently approved Moderna bivalent mRNA vaccines based on a mixture of ancestral SARS-CoV-2 with BA.1 or BA.4/BA.5 have been shown to confer high levels of neutralizing antibodies against several Omicron subvariants in clinical and animal studies [17, 18]. Thus, from the above studies and this study, bivalent boosters based on the Omicron variant could be an answer to improve protection against the currently circulating Omicron subvariants. The limitation of the study is primarily the small sample size which resulted in a large range of confidence intervals and precluded the use of more robust statistical tests to assess the statistical significance. However, this pre-clinical study presented sufficient evidence to permit further clinical investigation of the Omicron-based bivalent subunit vaccine.

## Acknowledgements

We are grateful to the team members at TFBS Bioscience Incorporation for performing animal experiments. We also thank the RNAi core facility at Academia Sinica (Taipei, Taiwan) for performing the pseudovirus neutralization assays.

## Author Contributions

T.-Y. K., C. E. L., and C.C. coordinated the projects. T.-Y.K., C. E. L., and Y.-J. L. designed the study and experiments. C.-C. W. and W.-H. T. produced the antigens used in the study. T.-Y. K. and Y.-J. L. supervised the animal experiments and assays. M.-Y. L. and L. T.-C. L. drafted the manuscript. All authors reviewed and approved the final version of the manuscript.

## Competing Interests

The authors declare no competing interests.

## Notes

### Competing Interest Statement

The authors have declared no competing interest.

